# Global meta-analysis reveals overall benefits of silvopastoral systems for biodiversity

**DOI:** 10.1101/2023.07.30.551160

**Authors:** Ricardo Perez-Alvarez, Julián Chará, Lauren D. Snyder, Michelle Bonatti, Stefan Sieber, Emily A. Martin

## Abstract

Domestic livestock grazing accounts for roughly one quarter of the world’s terrestrial surface and is a leading driver of biodiversity loss. Yet, it also provides a critical livelihood for nearly one billion smallholder farmers, creating a paradox that highlights the need for conservation strategies to balance human and ecological needs. Silvopastoral systems (SPS) integrate trees with livestock pastures, offering a promising solution to boost livestock productivity while safeguarding natural areas and biodiversity. However, evidence for the biodiversity benefits provided by SPS is limited to studies focusing on specific geographic regions or taxa. Through a global meta-analysis of 45 studies spanning 15 countries, four biogeographic regions, and seven taxa, we provide the first quantitative synthesis evaluating how SPS affect biodiversity and community stability relative to treeless pastures and natural forests. Overall, we show that SPS harbor higher levels of biodiversity (i.e., richness, abundance, and diversity) and stability than treeless pastures, and perform comparably to nearby forests. However, variations exist across regions and taxa, with the strongest positive responses in tropical dry regions and for low-mobility taxa like invertebrates and plants. Mammals, birds, and soil microorganisms, on the other hand, showed no significant biodiversity differences between treeless pastures and SPS. Thus, integrating SPS and protected areas as complementary components of a multifunctional landscape will be key to halting multi-taxa biodiversity loss and building sustainable livestock systems. Our findings support the conservation potential of SPS, while underscoring the need for strategic implementation to maximize benefits for biodiversity conservation.

## Introduction

To meet the world’s future food security and sustainability needs, it is imperative to reduce the environmental footprint of livestock production. Livestock grazing accounts for 26% of the world’s terrestrial surface and is one of the main drivers of biodiversity loss, particularly in tropical regions (1). For instance, over three-quarters of deforested lands in the Amazon Basin have been converted to pastureland to raise cattle in extensive grazing systems (2). These pasture systems have detrimental effects on local ecosystem structure and function (3), and negative global impacts on biodiversity and carbon stocks (4). With rising incomes in the developing world, demand for animal products will continue to grow (5) and could further drive the expansion of pasturelands at the expense of native forests and other biodiversity-rich ecosystems. Yet, an estimated one billion small-holder farmers, mainly in developing countries, depend on livestock for food and income (1), making it crucial to implement conservation strategies that balance human and ecological needs in these human-modified landscapes.

Silvopastoral systems (SPS)—multifunctional agroforestry arrangements that combine trees with livestock pastures—offer an alternative to counteract prevalent extensive treeless ranching systems. Compared to treeless pastures, SPS provide several benefits, including improved soil fertility and forage quality (6), reduced soil erosion and water loss (7), and enhanced carbon sequestration and storage (8). SPS also tend to support higher biodiversity than treeless pastures by providing wildlife habitat and increasing connectivity between remnants of natural vegetation (9). Alongside these environmental benefits, ranchers are increasingly receptive to adopting SPS due to their potential to increase livestock productivity and strengthen rural livelihoods (10, 11). Hence, scaling up SPS holds promise for balancing livestock production, conservation, and sustainable landscape restoration. As SPS are increasingly promoted as a restoration strategy in degraded grazing landscapes (10), there is a need to better understand the conservation value of these systems to support biodiversity.

Although SPS tend to enhance species diversity relative to treeless pastures (11), there are discrepancies and knowledge gaps that make generalizations difficult. First, the magnitude and direction of biodiversity responses to SPS differ across and among taxonomic groups. For example, studies in the Argentine Chaco Region found no differences in bird and mammal diversity between SPS and tree-less pastures (12), but dung beetle diversity was higher in SPS (13). Even within the same taxonomic group, there can be variation between different biodiversity dimensions. For instance, changes in dung beetle species richness in response to SPS do not necessarily reflect total abundance or species diversity changes (14). Understanding how such discordant changes across taxa and metrics affect overall biodiversity responses to SPS would enable a more accurate assessment of their conservation value.

Second, geographical variability may also influence SPS impacts on biodiversity. While recent efforts to implement SPS have targeted South American countries, similar systems are widely practiced in the European Mediterranean region (e.g., Dehesas and Montados) and, to a lesser extent, in temperate areas (15, 16). Indeed, SPS are currently deployed on 550 million hectares of land worldwide (17), representing biogeographic regions with different native biodiversity, climatic regimens, and ecosystem properties. Several studies have documented geographic variation in the biodiversity responses to SPS, and climatic factors such as temperature and precipitation have been proposed as the major drivers of these geographic trends (13, 17). In general, SPS can be more effective in recovering biodiversity relative to treeless pastures in regions dominated by tropical moist forests compared to dry forests or savannas (13, 18). These findings, however, stem from localized case studies lacking sufficient details to generalize outcomes confidently. Thus, determining the extent to which biodiversity responses to SPS differ across a wide range of latitudes, and the driving factors behind these variations remain important knowledge gaps.

Third, there is no standard approach to quantify the effectiveness of SPS in maintaining biodiversity. The ecological performance of agroforestry practices is often assessed by comparing mean levels of biodiversity between the agroforestry system and a baseline low-diversity or intensive agricultural state (e.g., treeless pastures) (19). While this approach provides a measure of how different agricultural practices perform compared to one another, it does not provide a picture of how well agroforestry schemes perform relative to natural systems. Conclusions about the conservation benefits of SPS could change if the baseline comparison were natural reforestation efforts or primary forests, rather than agriculture systems (20). Indeed, some agroforestry practices have positive short-term biodiversity responses relative to intensive agriculture, but have limited success as a conservation strategy compared to a reference natural ecosystem (21). Appropriate baselines are therefore fundamental to assessing whether SPS practices can restore the functioning of degraded pastures to native habitat levels, and serve as a tractable alternative to more intensive forms of agriculture.

In addition to a lack of appropriate baselines, current assessments of SPS do not provide a complete picture of biodiversity impacts. Converting forests to treeless cattle pastures, for example, could affect not only average biodiversity, but also community stability (i.e., temporal constancy in the aggregated abundance of all species in a community) (22). Understanding changes in community stability is critical as they can have profound effects on multiple ecosystem functions and the services they underpin, such as plant productivity, pollination, and pest control (23). While there is empirical evidence that community stability in treeless pasture systems is compromised compared to natural systems (24), the relationship between SPS and community stability remains to be explored. SPS are hypothesized to support more stable communities than treeless pastures by providing structurally complex environments whose microclimatic conditions and vegetation structure resemble those of forests and woodlands (18). Yet, whether this hypothesis is empirically supported across global biomes is unknown.

Here, we aim to disentangle the range of parameters that could influence the ability of SPS to support biodiversity and community stability by performing a global synthesis across a broad spectrum of taxa, geographies, and reference systems. We conducted an extensive literature review followed by a meta-analysis to ask: (1) Do SPS influence biodiversity and community stability relative to treeless pastures and natural forests? (2) If so, to what extent are these effects moderated by latitude and local climatic factors (i.e., precipitation and temperature)? (3) Are the strength and direction of these effects different across biogeographic regions and taxonomic groups? Our synthesis allows us to draw empirical generalities about the importance of SPS for biodiversity conservation and elucidates the conditions in which SPS could best support biodiversity.

## Results

Our meta-analysis was based on 45 studies that yielded 143 comparisons of biodiversity between SPS and treeless pastures, and 103 comparisons between SPS and natural habitats. The most examined biodiversity metric was species abundance (64%), followed by richness (30%) and diversity (6%). These primary studies were conducted in 15 countries representing four geographic regions: tropics (72%), subtropics (13%), Mediterranean (6%), and temperate (9%) (Fig. 1A). Insects (37%), macroinvertebrates (19%), and birds (17%) were the most prevalent taxonomic group used in comparisons. Soil bacteria and fungi (7%), plants (5%), and mammals (3%) were less represented (Fig. 1B).

**Figure 1.**
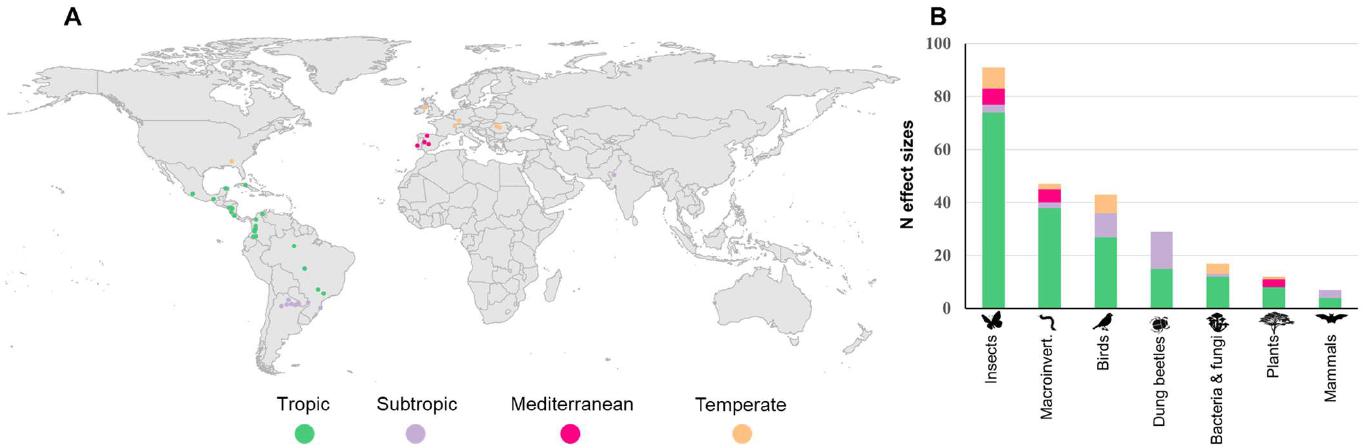
The geographical and taxonomic distribution of the studies included in the meta-analysis. The meta-analysis comprises 246 effect sizes from 45 independent studies. (**A**) Global distribution of studies included in the meta-analysis, with colors representing different biogeographic regions. (**B**) Relative distribution of studied taxonomic groups across biogeographic regions.

### Effects of SPS on biodiversity and community stability

SPS had significantly higher overall biodiversity than treeless pastures (mean Hedges’ g = 1.12, 95% confidence interval (hereafter CI) = 0.62 to 1.62, n = 143; Fig. 2). This pattern was consistent across all metrics, including species richness (g = 1.30, CI = 0.65 to 1.96, n = 36), abundance (g = 0.92, CI = 0.20 to 1.64, n = 93), diversity (g = 1.00, CI = 0.12 to 1.87, n = 7), and marginally for community stability (g = 0.29, CI = -0.02 to 0.60, n = 93). In contrast, we found no evidence that biodiversity differed between SPS and natural habitats for any of the biodiversity metrics assessed (overall: g = -0.12, CI = -0.59 to 0.35, n = 103; richness: g = - 0.29, CI = -0.83 to 0.26, n = 38; abundance: g = 0.25, CI = - 0.35 to 0.84, n = 47; diversity: g = 0.68, CI = -0.41 to 1.77, n = 8; community stability: g = 0.05, CI = -0.13 to 0.21, n = 47; Fig. 2). Sensitivity analysis confirmed the robustness of these results to geographic, taxonomic, and publication bias (tables S4 to S6 and fig. S2).

**Figure 2.**
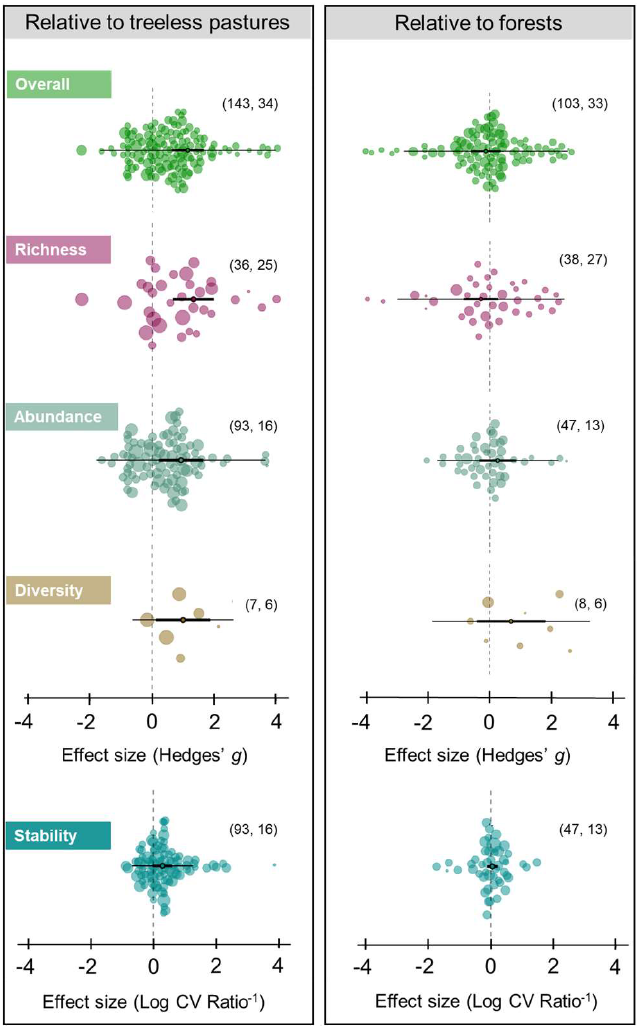
Effects of silvopastoral systems (SPS) on biodiversity and community stability relative to treeless pastures and forests. Points represent mean Hedges’g values, thick bars 95% confidence intervals (CI), and thin bars 95% prediction intervals. Each colored background point is an effect size, and its diameter is scaled by the precision of that estimate (1/SE); the larger the diameter, the more precise. A positive effect size means biodiversity was higher in SPS than in the corresponding control. Effect sizes are considered significant when the 95% CI does not cross zero (vertical dashed line). The number of effect sizes and studies included in each metric are displayed in parentheses, respectively.

Despite overall positive biodiversity responses to SPS, most models (57%) had a large and significant amount of total heterogeneity (tables S1 to S3), mainly attributed to between-study heterogeneity (I^2^ > 0.65). Subgroup analysis by biogeographic region and taxon showed that independently neither of them significantly explained this variability, but their interaction did (SPS relative to treeless pastures: omnibus F-test, F_19,123_ = 2.117, P = 0.008; SPS relative to forests: omnibus F-test, F_15,87_= 0.990, P = 0.0473).

We could not fully explore the effect of this interaction in our analysis due to the unbalanced distribution of taxa across regions and the high number of combinations (7 taxa groups x 4 biogeographic regions = 28 combination levels, Fig. 3). Nevertheless, similar analyses conducted with subsets of individual taxa or regions revealed important patterns in the direction and magnitude of biodiversity trends. Specifically, across regions, SPS supported higher biodiversity of insects, dung beetles, macroinvertebrates (other than insects), and plants relative to treeless pastures, whereas birds, mammals, and bacteria and fungi showed no response (Fig. 4A and Fig. S1). When comparing SPS to natural habitats, only dung beetles showed a significant decline in SPS, whereas all other taxa showed no significant differences (Fig. 4B and Fig. S1). Similarly, we found differences in the biodiversity responses to SPS between biogeographic regions. SPS showed higher overall biodiversity than treeless pastures in tropical and subtropical regions, whereas SPS and treeless pastures performed comparably in Mediterranean and temperate regions (Fig. 4C). Notably, SPS retained the same overall biodiversity levels as natural habitats in all biogeographic regions (Fig. 4D).

**Figure 3.**
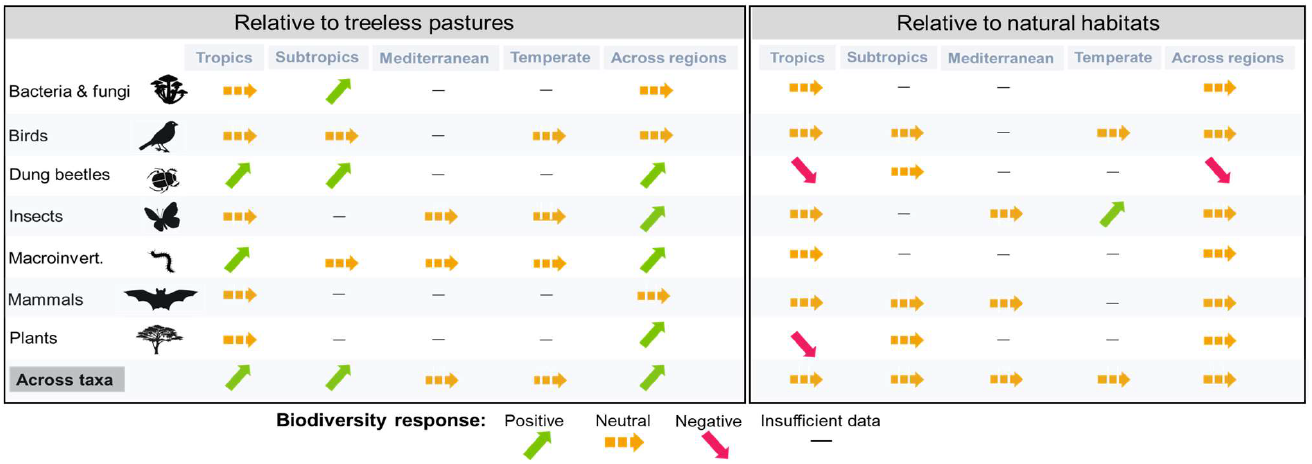
Summary of effects of silvopastoral systems (SPS) on biodiversity across taxa and biogeographic regions. For statistics, see Fig. 4 and supplementary tables S1-S3.

**Figure 4.**
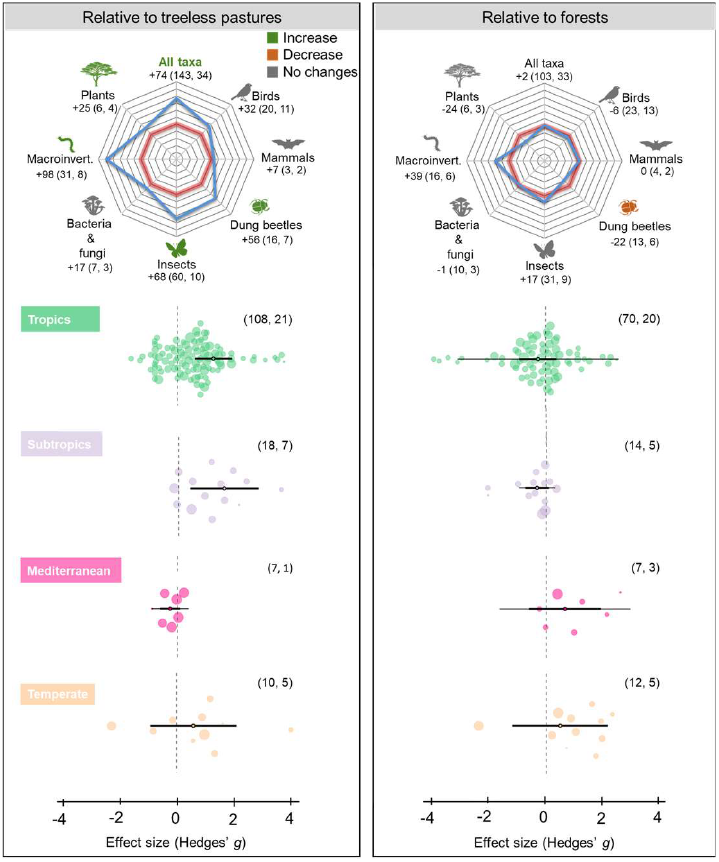
Effects of silvopastoral systems (SPS) on biodiversity across taxa and biogeographic regions. (**A** and **B**) Mean percentage change in biodiversity (i.e., species richness, abundance, or diversity) in SPS relative to treeless pastures and forests. In radar charts, red rings represent the baseline (i.e., no change in biodiversity with respect to the reference system), and blue rings represent a deviation away from the baseline; each interval corresponds to a 20% increase (from the red ring towards the outside edge) or decrease (from the red ring towards the center) in biodiversity. Mean values for changes in biodiversity are displayed below each taxa silhouette. Values were calculated using log response‐ratio effect sizes, which were back‐transformed and converted to percentage change. The number of effect sizes and studies included for each taxon are displayed in parentheses, respectively. (**C** and **D**) Relative effects of SPS on biodiversity across biogeographic regions. Points represent mean Hedges’g values, thick bars 95% confidence intervals (CI), and thin bars represent 95% prediction intervals. Each background point is an effect size, and its diameter is scaled by the precision of that estimate (1/SE); the larger the diameter, the more precise. A positive effect size means biodiversity was higher in SPS than in the corresponding control. Effect sizes are considered significant when the 95% CI does not cross zero (dashed line). The number of effect sizes and studies included in each geographic region are displayed in parentheses, respectively.

### Factors explaining the relative performance of SPS on biodiversity and community stability

Mediterranean and temperate regions were excluded from this analysis due to data paucity (n < 3 for each region). In general, latitude and precipitation significantly influenced the relative performance of SPS on biodiversity and community stability, whereas temperature had no influence on any of the metrics examined (Fig. 5 A and B). However, the relative importance of each factor varied depending on the biodiversity metric and reference system.

**Figure 5.**
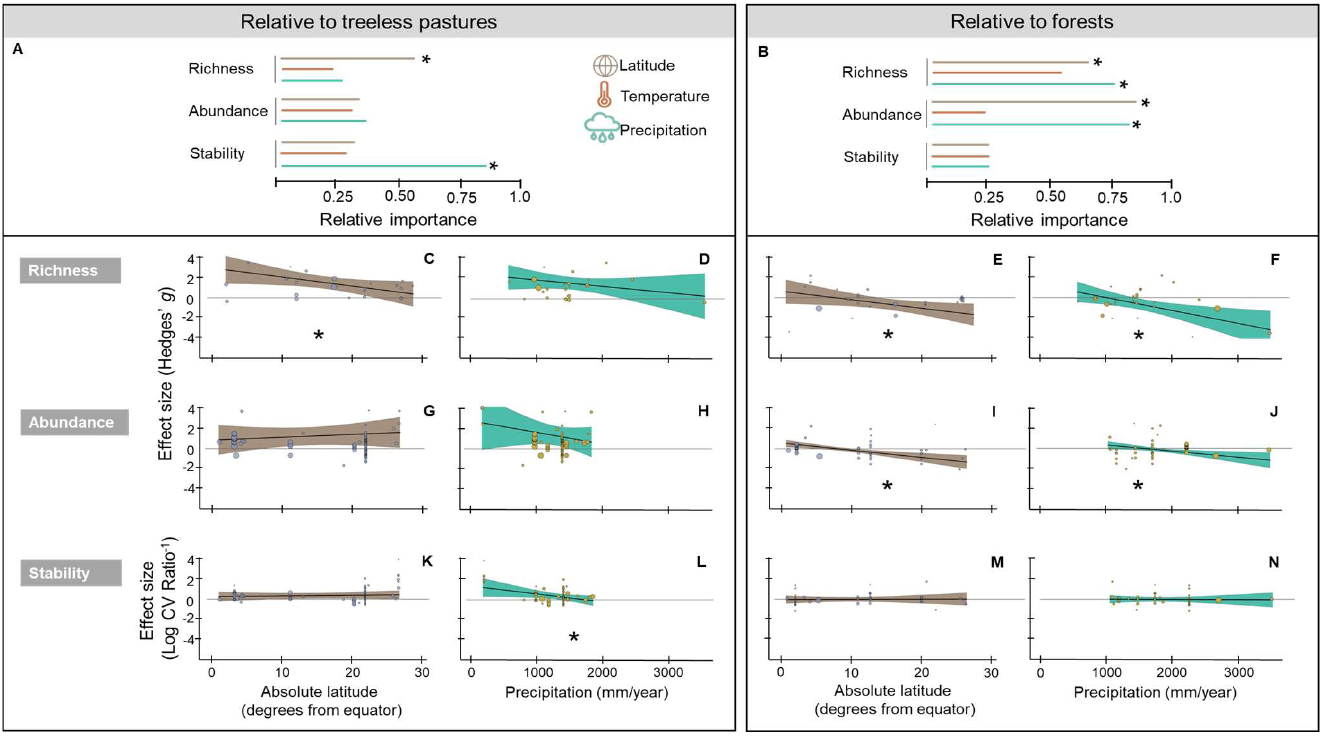
Relative importance of latitude, temperature, and precipitation in explaining the biodiversity performance of silvopastoral systems (SPS) when compared to treeless pastures and forests. (**A** and **B**) Variable importance ranges between 0 and 1; the higher the value, the more important the moderator factor. Asterisks indicate statistically significant variables (omnibus tests, P < 0.05). (**C** to **N**) Meta-regression models showing the moderating effects of precipitation and latitude on the performance of SPS relative to treeless pastures and forests. Lines depict predicted trends from meta-regression models and associated 95% confidence intervals (shaded areas). Each background point is an effect size, and its diameter is scaled by the precision of that estimate (1/SE); the larger the diameter the more precise. A positive effect size means that a given biodiversity metric was higher in SPS than in the corresponding control. Asterisks represent statistically significant results (omnibus tests, P < 0.05).

For models explaining variation in species richness and community stability between SPS and treeless pastures, our analysis showed that latitude and precipitation were the most important predictors (richness: Fig. 5 C and D, stability: Fig. 5 K and L). Increasing latitude and precipitation reduced SPS performance relative to treeless pastures. These findings suggest that SPS supported greater species richness and stability (relative to treeless pastures) in drier regions (mean annual precipitation < 2000 mm) of the tropics (i.e., lower latitudes). In line with these findings, we also found that SPS implemented in drier tropical regions more effectively maintained levels of richness and abundance, comparable to those in natural forests (richness: Fig. 5 E and F, abundance: Fig. 5 I and J). No significant predictors were found for species abundance in SPS compared to treeless pastures (Fig 5. G and H) and community stability in SPS relative to forests (Fig. 5 M and N). There were insufficient data to conduct the analysis on species diversity.

## Discussion

Silvopastoral systems (SPS) are increasingly seen as a win-win agricultural strategy that can meet the rising demand for animal products while also preserving remaining natural areas and biodiversity (6, 9). To better understand the conservation value of SPS, we conducted a global metaanalysis of 45 studies to examine their impact on biodiversity across taxa, biogeographic regions, and biodiversity metrics. Our results revealed that SPS have an overall positive effect on biodiversity and community stability compared to treeless pastures. We further found that SPS can, on average, support levels of biodiversity and stability comparable to those of nearby forests. However, these effects were highly variable, indicating that the range of biodiversity outcomes of SPS interventions varies widely. Differences across taxa, biogeographic regions, and local climatic factors explain part of this heterogeneity. Below, we discuss these sources of variability, outline their implications for biodiversity conservation, and highlight key remaining knowledge gaps.

While the overall effect of SPS on biodiversity, relative to treeless pastures, was positive, our analyses revealed several important differences across taxonomic groups. This variability might reflect differences among taxa in key ecological traits—such as mobility and dispersal abilities — that mediate biodiversity responses. For example, previous work suggests that plants and arthropods with limited mobility have a stronger response to on-farm conservation activities than to landscape factors, while taxa with greater dispersal abilities can exhibit the opposite response (25). Accordingly, we found that plants and invertebrates (i.e., insects, dung beetles, and macroinvertebrates) were more diverse in SPS than in treeless pastures, while mammals and birds did not present differences between treeless pastures and SPS. Greater mobility may enable birds and mammals to capture resources beyond an agricultural pasture, thus buffering them against the adverse effects of local agricultural intensification (i.e., treeless pastures). It follows that for taxa with larger foraging ranges, SPS may be insufficient to maintain stable populations unless surrounding areas of natural habitat are also protected (26). In addition to taxa with greater mobility, we also found that soil microorganisms (i.e., bacteria and fungi) did not vary between SPS and treeless pastures. The lack of positive effects of SPS on soil microorganisms is unexpected and may indicate that soil type and local management are more important than tree cover for those organisms, as previously reported (27). Potential interaction between SPS and soil microorganisms, however, deserves more attention as our inferences are based on a small sample size (i.e., n = 7).

Our analyses clearly demonstrate that different organisms exhibit contrasting responses to SPS, suggesting we must be cautious in applying single taxa measures as a proxy for broader biodiversity responses to SPS. Instead, multiple functional groups should be evaluated if we are to provide robust recommendations about the value of SPS for safeguarding biodiversity. Furthermore, while the local implementation of SPS may be sufficient for preserving some taxa, a broader conservation agenda that integrates both local and landscape strategies is needed to restore biodiversity across taxa.

In addition to contrasting responses across taxa, our results show that biodiversity responses to SPS varied across biogeographic regions. Implementing SPS in tropical and subtropical regions had an overall positive effect on biodiversity compared to treeless pastures. Conversely, in Mediterranean and temperate regions, there was little to no difference in biodiversity between SPS and treeless pastures. Geographic variation in species’ sensitivity to land use and local climatic changes is one possible explanation for these differences. Indeed, previous studies indicate that tropical species are often more sensitive to land-use change than their temperate counterparts (28). This effect is largely due to a long history of agriculture in temperate regions, which likely already filtered out the most sensitive species even from natural habitats, rendering the remaining communities less sensitive to contemporary disturbances (29). Species inhabiting areas with climatic seasonality also tend to be more tolerant to local climatic changes following tree cover loss (30). In contrast, tropical and subtropical biomes are more sensitive to current land-use change as they are characterized by a shorter history of intensive agriculture, a lower degree of seasonality, and biota with smaller range sizes (28). Our meta-analysis reinforces earlier conclusions regarding the importance of agroforestry as a strategy for reducing the vulnerability of tropical and subtropical biodiversity to global drivers of land use change (31). Although we found little to no effect of SPS on biodiversity in temperate and Mediterranean regions, we are cautious in giving too much weight to this result, as it is based on small sample sizes (i.e., temperate: n = 10; Mediterranean: n = 7). Nevertheless, this geographic variation implies that climatic and historical land use conditions could impact the performance of SPS and warrants further investigation.

In line with these results, we found that the effect of SPS on biodiversity was stronger in drier tropical regions compared to wetter regions. Just as with the differences across biomes, variation in the sensitivity of biodiversity to land use changes could explain the stronger positive responses to SPS in drier regions. Communities in dry climates (defined here as regions with mean annual precipitation lower than 2000 mm) are particularly vulnerable to the loss of vegetation and forest cover, as they are already facing physiological limitations due to low water availability (32). The tree canopy layer created in SPS can provide thermal and humidity refuges that protect biodiversity in arid and semiarid environments. In fact, SPS can reduce mean annual temperatures by 2-3°C and increase relative humidity by 10-20% compared to treeless pastures (33). This has important implications for the conservation and management of tropical dry forests, as they are both highly threatened by agricultural pressures (34) and are particularly vulnerable to projected increases in drought frequency and severity associated with climate change (35).

Despite substantial variation across taxa and regions, overall, SPS supported higher species richness, abundance, and diversity than treeless pastures. This result reinforces previous findings indicating that integrating pastures with trees can create more habitats and resources—such as food supply, nesting, or breeding sites—that better support biodiversity than treeless pastures (36, 37). In addition to highlighting the potential role of SPS in preserving farmland biodiversity, we demonstrate that SPS can enhance community stability. SPS can confer greater stability by providing habitat for a diversity of species that occupy similar functional niches (functional redundancy) (38) or that are functionally similar, but respond differentially to environmental change (response diversity) (39). Although not tested here, previous empirical evidence indicates that greater community stability in SPS would lead to a more stable provision of ecosystem services such as food production, carbon sequestration, and soil fertility (40–42).

Importantly, our findings revealed that SPS harbored levels of species richness, abundance, diversity, and community stability comparable to surrounding natural forests. This finding was consistent across biogeographic regions and taxa—albeit the one exception was dung beetles which showed lower diversity in SPS than in natural forests. This similarity between SPS and natural forests reinforces the potential of SPS to serve as a tool for biodiversity conservation. This finding, however, has its limitations. It is worth noting that our analysis was based on metrics of species richness, abundance, and diversity that do not capture subtler effects on species identity (43). It could be argued that the presence of forest specialists, or endangered or endemic species is more informative for conservation decisions than taxonomic metrics alone (44).

Consequently, the lack of biodiversity differences between SPS and natural forests does not mean they are ecologically equivalent. Indeed, a previous study showed that similar bird species richness between natural forests and SPS belied marked differences in bird community composition (45). Similarly, SPS may host mammal assemblages with a higher proportion of generalist species and fewer forest specialists compared to natural forests (36). Forest remnants seem then fundamental to maintaining viable populations of many species, especially for forest specialists and area-demanding species (26). Further research is needed, however, to establish the exact role that SPS can play in supporting landscape conservation efforts by providing secondary habitats to disturbance-tolerant forest species and increasing connectivity between forest fragments (e.g., (46)). This is particularly relevant in tropical forests as they are becoming increasingly fragmented by the widespread and rapid expansion and intensification of agricultural activities (47). Focusing on the response of forest specialists and species composition may therefore provide further insights into the conservation value of SPS relative to natural forests.

### Limitations and knowledge gaps

Our meta-analysis is subject to limitations inherently associated with synthesizing data on a global scale, which should be kept in mind when interpreting our results. First, our dataset was unevenly distributed across biogeographic regions and taxonomic groups. More than a third of the data on the effects of SPS on biodiversity come from tropical and subtropical regions in Central and South America, and are primarily from studies focused on invertebrates and birds (i.e., 85% of the overall studies). Our sensitivity analysis revealed that our conclusions are relatively robust to geographic and taxonomic bias. Yet, future studies on the effects of SPS on understudied taxa (e.g., mammals, plants, and soil microorganisms), especially in temperate and Mediterranean regions, will be essential to enable a more comprehensive view of biodiversity responses to SPS. Second, our conclusions are based on taxonomic metrics— richness, abundance, and diversity—because they are the most widely used measures to evaluate SPS effectiveness. However, as discussed above, these metrics do not capture the many dimensions of biodiversity (48). Future studies should include a broader suite of biodiversity metrics (e.g., similarity indices, phylogenetic diversity, functional diversity). Similarly, looking beyond taxonomic metrics to focus on how these biodiversity changes influence ecosystem functions and services is a challenge that is yet to be addressed at a global scale (but see (49)). Third, aggregated biodiversity responses for a given taxon do not always reflect species-specific responses within that group. For instance, as noted by past work, arthropods span such a diverse taxon that individual species may display divergent responses to SPS (50). Future research should therefore explore SPS impacts at a finer taxonomic resolution than we have here (e.g., Family level). Fourth, SPS arrangements vary widely both within and across regions. These configurations can range from scattered eucalyptus or pine trees planted among exotic grasses to dense native tree plantations and native pastures. It is likely that species respond differently to this variability, but this premise has yet to be confirmed due to the shortage of studies that provide comprehensive information about tree diversity and its spatial arrangement in SPS (e.g., scattered trees, living fences, or densely mixed stands).

### Conclusions and policy implications

Despite these limitations, our study clearly shows that SPS is a potential pathway to restoring biodiversity in degraded landscapes. This biodiversity, in turn, could yield enormous benefits for carbon sequestration, human livelihoods, and many ecosystem services. Consequently, ambitious plans have been put forward in many forest-rich countries to restore more than five million hectares of degraded land through the widespread adoption of SPS (55, 56). Our results support the conservation potential of such initiatives by showing that SPS provide biodiversity benefits at the farm level. However, our study also demonstrates that the potential for SPS to support biodiversity varies widely depending on several factors related to historic land-use, geographic region, and climatic conditions. Such context-dependency indicates that we must move beyond simplistic definitions of targets based on how much land to restore and advance to prioritizing where and how to implement SPS to maximize benefits for both nature and people. Our results also show that although SPS represent a clear benefit to biodiversity compared to treeless pastures, they are not a silver bullet for conservation. In particular, SPS benefit plants and invertebrates most, whereas more mobile, landscape-dependent taxa benefit less. Thus, combining SPS and protected areas as complementary components of a multifunctional landscape will be key to minimizing biodiversity loss in livestock-dominated landscapes. Alongside this, appropriate safeguards are needed to ensure that policies promoting the widespread adoption of SPS will not incentivize further natural habitat degradation (e.g., deforestation spillovers, (55)). Mainstreaming SPS at the expense of native vegetation will be detrimental—not beneficial—to biodiversity (57). Finally, policy decisions aimed at reconciling conservation and smallholder production, such as upscaling SPS, need to address the local socioeconomic barriers hindering the adoption of sustainable agricultural practices (58). Identifying areas that align high potential for biodiversity benefits with socioeconomic feasibility will be essential to meet countries’ pledges to restore degraded pasturelands.

## Materials and Methods

### Data collection and extraction

We conducted a literature search following a hierarchical procedure. First, we used Web of Science, Scopus, and Scielo as search engines to identify published studies that provided data on the effect of silvopastoral systems on biodiversity from 1998 to 2021. The initial search string was: [ALL (bird* OR mammal* OR reptile* OR amphibia* OR arthropod* OR insect* OR fish* OR plant* OR vegetation* OR bacteria* OR ants OR fauna OR “soil fauna” OR “soil bacterial community” OR pollinators OR Scarabaeidae OR “dung beetles”) AND ALL (“silvopastoral systems” OR “SPS” OR “pastures with trees” OR “silvopastoral” OR “silvopasture” OR “sustainable cattle” OR “cattle grazing systems” OR “sustainable livestock” OR “cattle ranching” OR “livestock” OR “livestock systems” OR pasture* OR “open pastures” OR cattle) AND (effective* OR performance* OR assessment* OR evaluation* OR estimate* OR comparison* OR contrast*) AND (biodiversity* OR “species diversity” OR diversity OR richness OR “species richness” OR abundance OR “species abundance” OR conservation OR “biodiversity conservation”)]. In a second step, we used unstructured literature searches by examining the reference lists and citing articles of relevant studies and recent systematic reviews. Finally, we explored references in the grey literature using expert knowledge of available information and searches on other platforms (e.g., ProQuest Dissertation, EBSCOhost Open Dissertations, preprints online repositories).

The initial search resulted in a set of 7,789 publications from which we retained studies meeting the following selection criteria based on the Population, Intervention, Control, and Outcome (PICO) framework (59). According to this framework, studies had to: (1) investigate any species (plant, animal, bacteria, and fungi) (Population); (2) measure biodiversity responses in both silvopastoral systems and controls (treeless pastures or natural habitats), in a replicated study design for comparison (Intervention and Control); (3) investigate biodiversity in terms of species richness, species abundance (i.e., abundance of individual species), abundance per assemblage (i.e., when values are summed across groups of unidentified species), or species diversity as response variables (Outcome). Although biodiversity can be measured in various ways, we restricted our analysis to studies that calculated taxonomic-based measures of species richness, species abundance, and diversity indices, as they are commonly used in biodiversity studies. Studies that only measured functional or phylogenetic diversity were not included. Some studies included comparisons of more than one response variable (e.g., species richness and abundance) or various taxa. We considered each comparison as an independent observation and used the identity of the primary study as a random factor to control for potential correlations between comparisons within a primary study (see Data analysis). When researchers provided species diversity, richness, and abundance measures, we extracted only richness and abundance values to avoid pseudoreplication, as diversity indices are not calculated independently, but rather are derived from richness and abundance values. For studies reporting response variables at different time points, we kept only the last time point to capture long-term responses. If researchers tested different biodiversity treatments within SPS systems or controls, we followed the approach used in previous meta-analyses and extracted biodiversity measurements from the treatment with the highest biodiversity values (60).

Our final database included information from 45 studies (which yielded 246 effect sizes) published between 1998 and 2022 (see PRISMA flow diagram: fig. S1). From these studies, means, standard deviations, and sample sizes for SPS and controls were extracted from the text, tables, or figures using WebPlotDigitzer (61) and metaDigitise (62). If the information was not fully available in the main text or supplement information, we contacted the authors and requested the missing data (8 out of 26 responded positively). When we were unable to obtain this information, we excluded that article from further analysis. For the selected studies, we also collected information on the year of publication, location (latitude), biogeographic region, taxonomic group, and climatic conditions (temperature and precipitation). All retrieved data is available in the supplementary information (data file S1).

### Effect sizes

Standardized mean differences (Hedges’g) between biodiversity measures in SPS and its corresponding control site (treeless pastures or natural habitats) were used as effect sizes (63). Hedges’g was selected because it can account for small sample sizes and uneven variance between treatment groups (64). A positive effect size indicates higher biodiversity in SPS systems compared to controls. In parallel to Hedges’g, and to facilitate interpretation of the percentage of biodiversity change between groups, we also computed back-transformed log response ratios lnRR. To investigate the relative differences in community stability between SPS and controls, we calculated an additional effect size—the inverse of the logtransformed CV ratio (lnCVR^-1^)(65). The lnCVR-1 is analogous to the inverse of the coefficient of variation (CV−1) used in ecological fields to quantify stability (66). As with Hedges’g, a positive lnCVR^-1^ indicates higher community stability in SPS relative to controls.

### Data analysis

We analyzed variation in effect sizes for overall biodiversity (i.e., global effects across all biodiversity metrics) and separately for each response variable (richness, abundance, diversity, and community stability). Biodiversity differences between treatment groups were considered statistically significant if the 95% CI of the mean overall effect did not overlap with zero (67). To assess whether results differed across taxa and biogeographic regions, we conducted a subgroup analysis. For this, we clustered biodiversity measures into six taxonomic groups: birds, dung beetles, insects (other than dung beetles), mammals, plants, soil macroinvertebrates (excluding insects), and soil bacteria and fungi. The taxonomic categories we employed followed operational necessities to balance the level of taxonomic resolution (i.e., avoid a resolution that is too coarse to detect patterns) and robustness (i.e., categories with enough sample size). For instance, we found studies conducted on specific insect groups such as ants and bees, but their sample sizes were too small for robust separate comparisons (n < 10) (68). We also categorized studies into four biogeographic regions: tropics (< 23.5° latitude in both hemispheres), subtropics (between 20° and 40° latitude in both hemispheres), temperate (between 40° and 66.5° latitude in both hemispheres), and Mediterranean (spanning coastal and interior portions of the Mediterranean Basin). Subgroup analyses for taxonomic group, biogeographic regions, and their interactions were conducted for overall biodiversity measures (i.e., global effects across all biodiversity metrics) as insufficient data were available for robust comparisons separately on species richness, abundance, diversity, and community stability.

Multilevel linear mixed-effects models (‘rma.mv’) with a restricted maximum-likelihood estimator (REML) were used to estimate mean overall differences between treatment groups. To account for the non-independence of multiple effect sizes within the same study, all models were fitted using case number nested within study ID as a random effect (69). These models account for both sampling variance within studies and between-study variance. Heterogeneity was tested with a Q test and additionally represented by I2, the ratio of heterogeneity (i.e., between-study variability) to the total variability (i.e., the sum of between- and within-study variability).

To further explain the variation in effect sizes, we fitted linear mixed ‐ effects meta ‐ analysis models (i.e., ‘rma.mv’) using maximum likelihood estimation. Latitude, mean temperature, and mean precipitation were included as moderators. Study latitude was defined as the central point of the study region, measured as the distance from the equator in decimal degrees (i.e., absolute values of latitude spanned from 1 to 54°). Similarly, our mean temperature and precipitation values were calculated for the same locations, using data from the original papers and the WorldClim database when the data were not reported (70). We applied an information-theoretic approach to model selection and multi-model inference to determine the relative importance of each moderator variable (71). Due to collinearity among moderators and because it may hamper interpretation, we did not include interaction terms in our models.

To build our models, we first created a candidate set of models with all possible additive linear combinations of moderators and retained the highest-ranked models based on the corrected Akaike’s information criterion (AICc ≤ 2) (71). From the set of plausible models, we computed the model-averaged coefficients and the relative importance of each moderator as the sum of Akaike’s weights of all the selected models in which the moderator was included. Variable importance parameters range between 0 and 1; the higher the value, the more important the moderator factor. The significance of the moderators was determined with the Omnibus test, against a χ2-distribution. Lastly, with the set of significant moderators, we ran metaregression models to explain how individual moderators influence SPS performance relative to treeless pastures and forests.

### Publication bias and sensitivity analysis

Meta-analysis results may be affected by publication bias, i.e., the selective publication of articles finding significant effects over those which find non-significant effects.

Publication-bias analysis often includes Egger regression tests and funnel plots. However, meta-analytic datasets in ecology are not suitable for these methods due to high levels of heterogeneity and non-independent effect sizes. Following Nawasaka et al. (72), publication bias was tested by modifying the multi-level meta-analysis model to include the square of the inverse of effective sample size as a moderator. Furthermore, we tested the effect of time-lag bias (i.e., when statistically significant effects are published more quickly than smaller or non-statistically significant effects) by also including the year of publication (mean-centered) as a moderator. We considered models to be biased if the slope estimate of any of the moderators differed significantly from zero. For biased models, we assessed the impact of publication bias by applying the trim-and-fill method (73). We first using a compound symmetric structure (‘struct=CS’) and a value of 0.5 for rho (74). Using the aggregate data, we then fitted the trim-and-fill method by evaluating the asymmetry in reported study outcomes and re-estimating the average global effect size. If the bias-adjusted overall effect size retains the same sign and significance, the results of the meta-analysis are considered robust to possible publication bias (69). Although publication bias was detected in some models (particularly on models comparing SPS to treeless pastures), the direction and significance of bias-adjusted estimates remained unchanged, indicating that the impact of publication bias did not affect our overall conclusions (tables S4 to S6).

As our database had a strong geographic bias towards tropical regions (see results), we tested the robustness of our results by re-running the analysis with two different datasets: (i) a subset that excluded data from tropical regions, and (ii) a subset that only included data from tropical regions. A similar bias was evident for taxa, with insects accounting for 37% of all effect sizes. We re-ran the analysis with two additional subsets that (iii) excluded insect data, and (iv) excluded taxa representing < 7% of all effect sizes (i.e., bacteria and fungi, plants, and mammals, see results). The analysis of the four datasets gave qualitatively similar results (fig. S2), confirming the robustness of our results to both geographic and taxonomic bias. All analysis and data processing were carried out in R v.4.0.4 using packages, ‘metafor’ v.3.0-2 (75), ‘MuMIn’ (76), and ‘orchard’ (77).

## Acknowledgments

We thank all of the authors who contributed their data to this study. We also thank I. Osorio Laguna for assistance with data extraction, and the Zoological Biodiversity Group at the Leibniz University of Hannover for comments and suggestions on earlier drafts of the manuscript. Funding: This work was supported by the Leibniz Centre for Agricultural Landscape Research (ZALF) (Auftrag ZALF 111450) to MB and SS. RPA was supported by a Fulbright-Minciencias fellowship. LDS received funding through a Leibniz University Hannover Flexible Funds Grant (project number: 51171322).

## Author contributions

RPA, JC, LDS, and EAM conceived the idea and designed the study. RPA and JC compiled the dataset. RPA analyzed the data and led the writing. MB and SS contributed with funding acquisition. All authors interpreted the results and edited the manuscript.

## Data and materials availability

All data needed to evaluate the conclusions in the paper are present in the paper and/or the Supplementary Materials. The dataset used in this study will be available in data file S1 pending scientific review and a completed material transfer agreement. Additional data related to this paper may be requested from the corresponding author.

## Supplementary Materials

### Other Supplementary Materials for this manuscript include the following

Data file S1

**Fig. S1.**
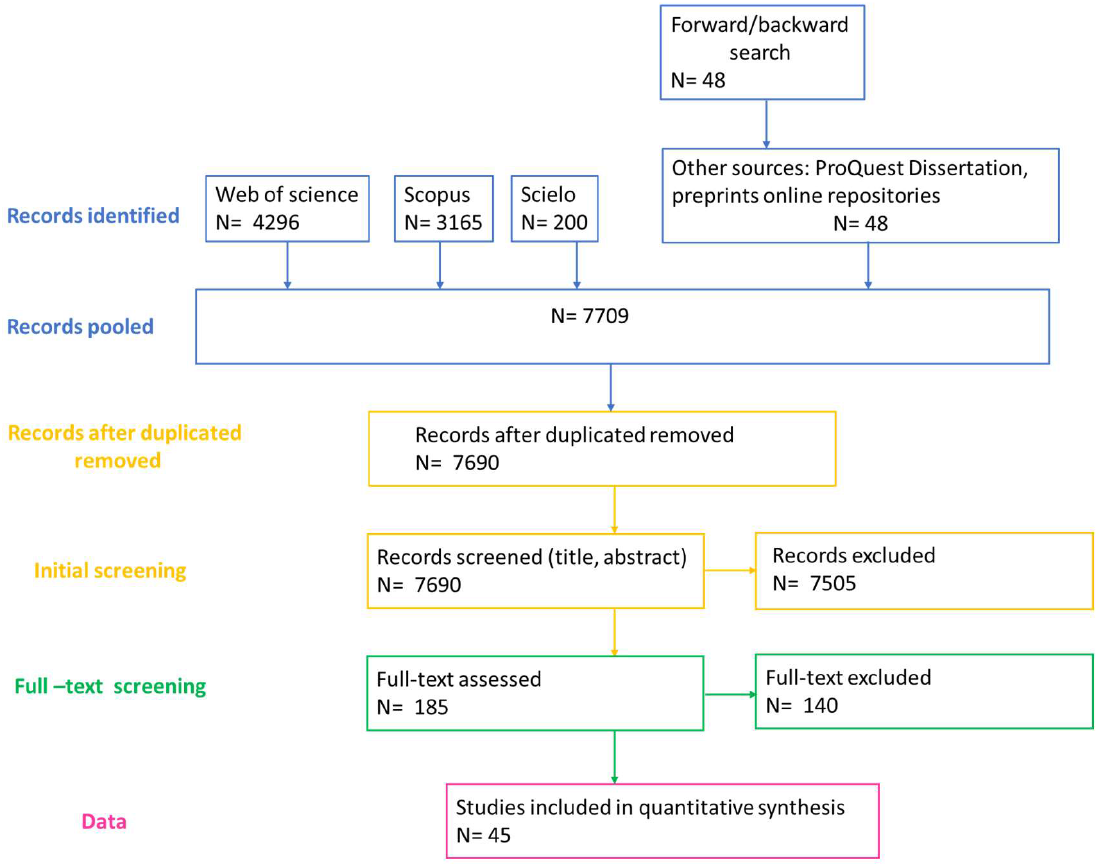
PRISMA flow diagram describing the process of searching, filtering and inclusion of studied used in our meta-analysis.

**Fig. S2.**
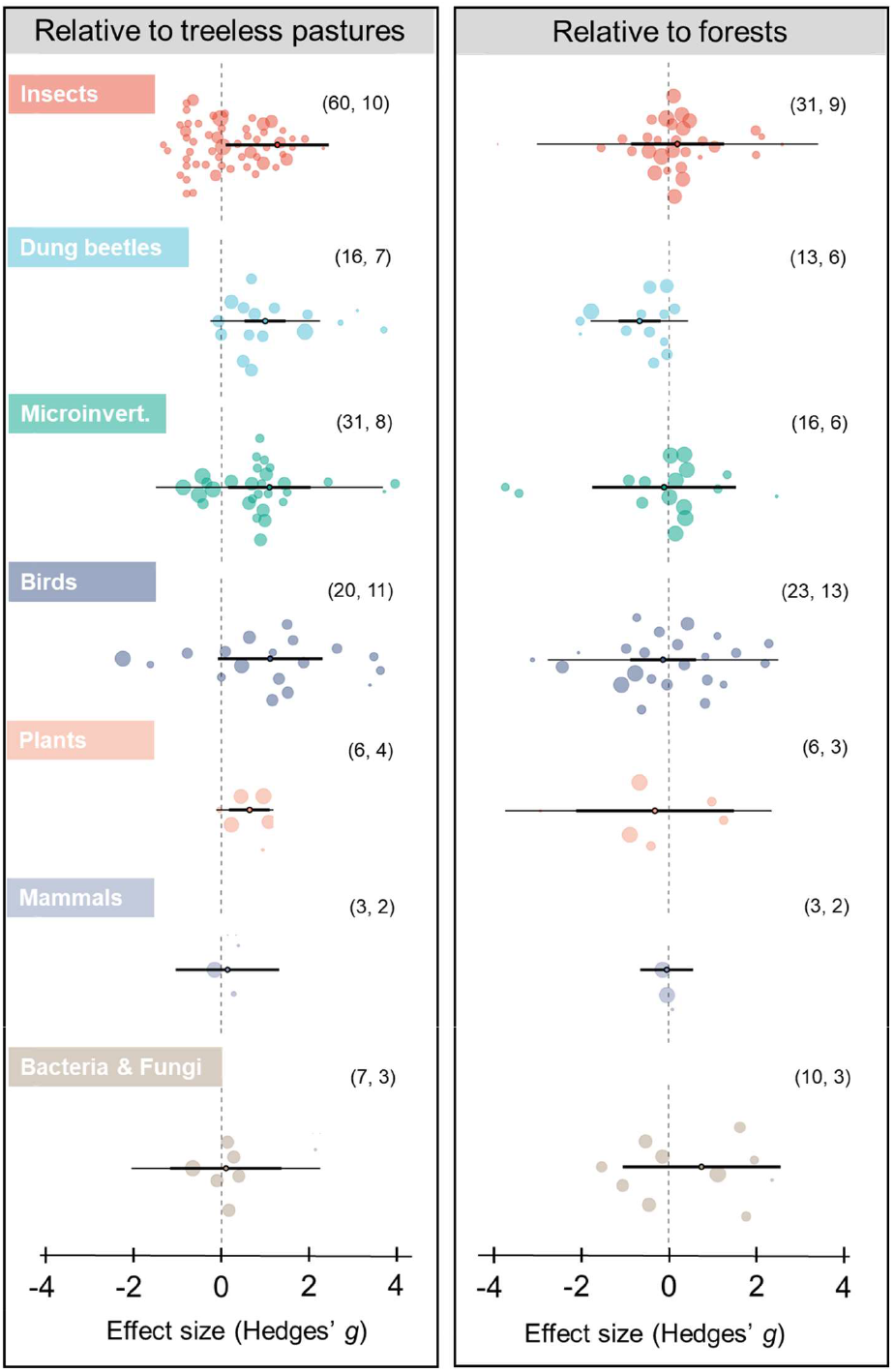
Relative effects of silvopastoral systems (SPS) on biodiversity across taxonomic groups. Points represent mean Hedges’*g* values, thick bars represent 95% confidence intervals (CI), and thin bars 95% prediction intervals. Each colored background point is an effect size, and its diameter is scaled by the precision of that estimate (1/SE); the larger the diameter, the more precise. A positive effect size means biodiversity was higher in SPS than in the corresponding control. Effect sizes are considered significant when the 95% CI does not cross zero (vertical dashed line). The number of effect sizes and studies included in each taxon are displayed in parentheses, respectively.

**Fig. S3.**
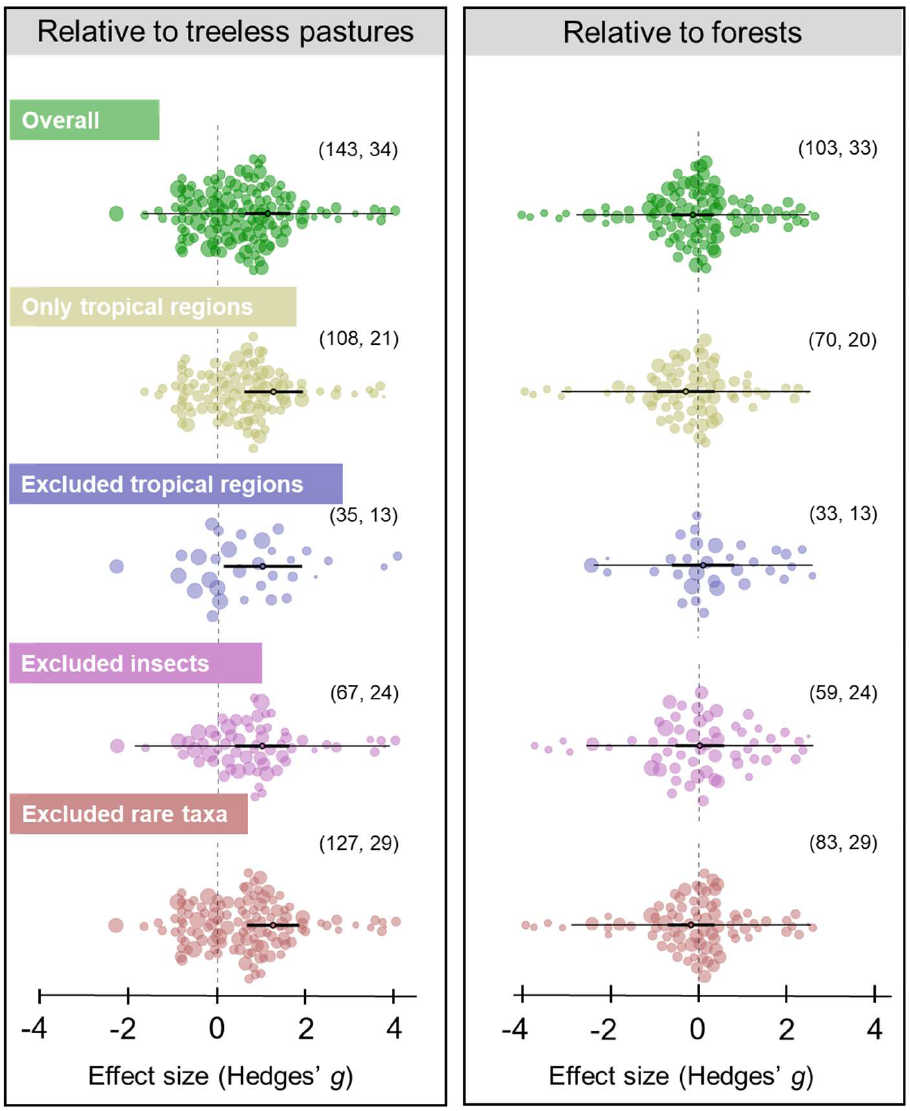
Sensitivity analysis confirmed the robustness of the effects of SPS on biodiversity to geographic and taxonomic bias. As our database was biased towards studies with insects in tropical regions, we re-ran the analysis with four different datasets to test the robustness of our results: (i) a subset that included only data from tropical regions, (ii) a subset that excluded data from tropical regions, (iii) a subset that included only data from insects, and (iv) a subset that excluded taxa representing < 7% of all effect sizes (i.e., bacteria and fungi, plants, and mammals). The points on the graph represent mean Hedges’ *g* values, the thick bars represent 95% confidence intervals (CI), and the thin bars represent 95% prediction intervals. Each colored background point represents an effect size, with the diameter scaled by the precision of the estimate (1/SE); larger diameters indicate more precise estimates. A positive effect size indicates that biodiversity was higher in SPS than in the corresponding control. Effect sizes are considered significant when the 95% CI does not cross zero (vertical dashed line). The number of effect sizes and studies included in each subset is displayed in parentheses.

**Table S1.**
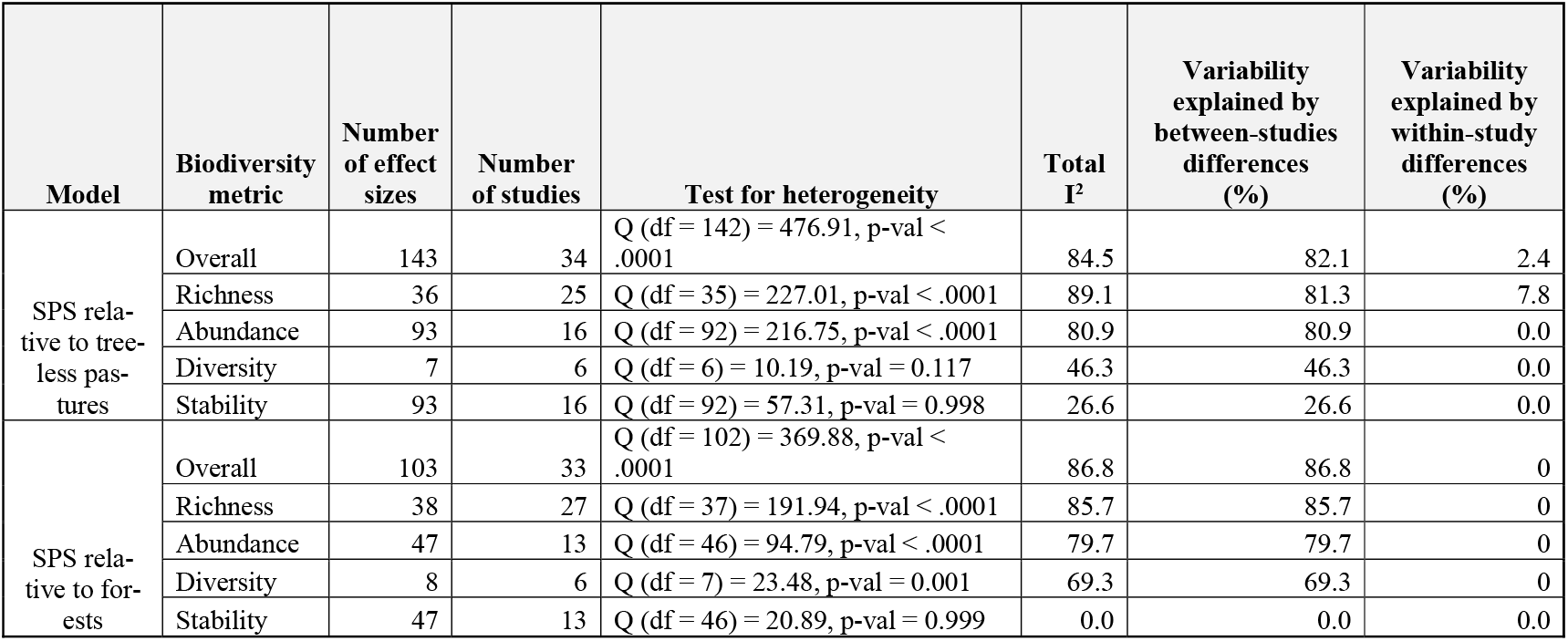
Assessment and quantification of heterogeneity in meta-analytic models across biodiversity metrics. A Cochran’s heterogeneity Q-test (*78*) evaluates whether the observed effect sizes’ variability is greater than what would be expected from sampling variability alone. A significant test result indicates that true effect size differences exist between studies in our data. The I^2^ statistic shows the percentage of variation across studies that is due to heterogeneity, rather than chance (i.e., sampling error or within study variability) (*79*). I^2^ values greater than 75% indicate substantial heterogeneity (*80*).

| Model | Biodiversity metric | Number of effect sizes | Number of studies | Test for heterogeneity | Total $I^2$ | Variability explained by between-studies differences (%) | Variability explained by within-study differences (%) |
| --- | --- | --- | --- | --- | --- | --- | --- |
| SPS relative to treeless pastures | Overall | 143 | 34 | Q (df = 142) = 476.91, p-val < .0001 | 84.5 | 82.1 | 2.4 |
|  | Richness | 36 | 25 | Q (df = 35) = 227.01, p-val < .0001 | 89.1 | 81.3 | 7.8 |
|  | Abundance | 93 | 16 | Q (df = 92) = 216.75, p-val < .0001 | 80.9 | 80.9 | 0.0 |
|  | Diversity | 7 | 6 | Q (df = 6) = 10.19, p-val = 0.117 | 46.3 | 46.3 | 0.0 |
|  | Stability | 93 | 16 | Q (df = 92) = 57.31, p-val = 0.998 | 26.6 | 26.6 | 0.0 |
| SPS relative to forests | Overall | 103 | 33 | Q (df = 102) = 369.88, p-val < .0001 | 86.8 | 86.8 | 0 |
|  | Richness | 38 | 27 | Q (df = 37) = 191.94, p-val < .0001 | 85.7 | 85.7 | 0 |
|  | Abundance | 47 | 13 | Q (df = 46) = 94.79, p-val < .0001 | 79.7 | 79.7 | 0 |
|  | Diversity | 8 | 6 | Q (df = 7) = 23.48, p-val = 0.001 | 69.3 | 69.3 | 0 |
|  | Stability | 47 | 13 | Q (df = 46) = 20.89, p-val = 0.999 | 0.0 | 0.0 | 0.0 |

**Table S2.** Assessment and quantification of heterogeneity in meta-analytic models across biogeographic regions. A Cochran’s heterogeneity Q-test (*78*) evaluates whether the observed effect sizes’ variability is greater than what would be expected from sampling variability alone. A significant test result indicates that true effect size differences exist between studies in our data. The I^2^ statistic shows the percentage of variation across studies that is due to heterogeneity, rather than chance (i.e., sampling error or within study variability) (*79*). I^2^ values greater than 75% indicate substantial heterogeneity (*80*).

| Model | Biogeographic region | Number of effect sizes | Number of studies | Test for heterogeneity | Total $I^2$ | Variability explained by differences among studies (%) | Variability explained by differences within study (%) |
| --- | --- | --- | --- | --- | --- | --- | --- |
| SPS relative to treeless pastures | Tropics | 108 | 21 | $Q (df = 107) = 247.57, p\text{-val} < .0001$ | 82.2 | 82.2 | 0.0 |
| | Subtropics | 18 | 7 | $Q (df = 17) = 71.18, p\text{-val} < .0001$ | 80.4 | 61.6 | 18.7 |
| | Mediterranean | 7 | 1 | $Q (df = 6) = 9.17, p\text{-val} = 0.164$ | 34.2 | 0.0 | 34.2 |
| | Temperate | 10 | 5 | $Q (df = 9) = 89.19, p\text{-val} < .0001$ | 90.9 | 39.8 | 51.2 |
| SPS relative to forests | Tropics | 70 | 20 | $Q (df = 69) = 224.87, p\text{-val} < .0001$ | 89.1 | 89.1 | 0.0 |
| | Subtropics | 14 | 5 | $Q (df = 13) = 13.20, p\text{-val} = 0.433$ | 15.0 | 15.0 | 0.0 |
| | Mediterranean | 7 | 3 | $Q (df = 6) = 12.16, p\text{-val} = 0.059$ | 59.7 | 59.7 | 0.0 |
| | Temperate | 12 | 5 | $Q (df = 11) = 89.92, p\text{-val} < .0001$ | 90.1 | 86.8 | 3.3 |

**Table S3.**
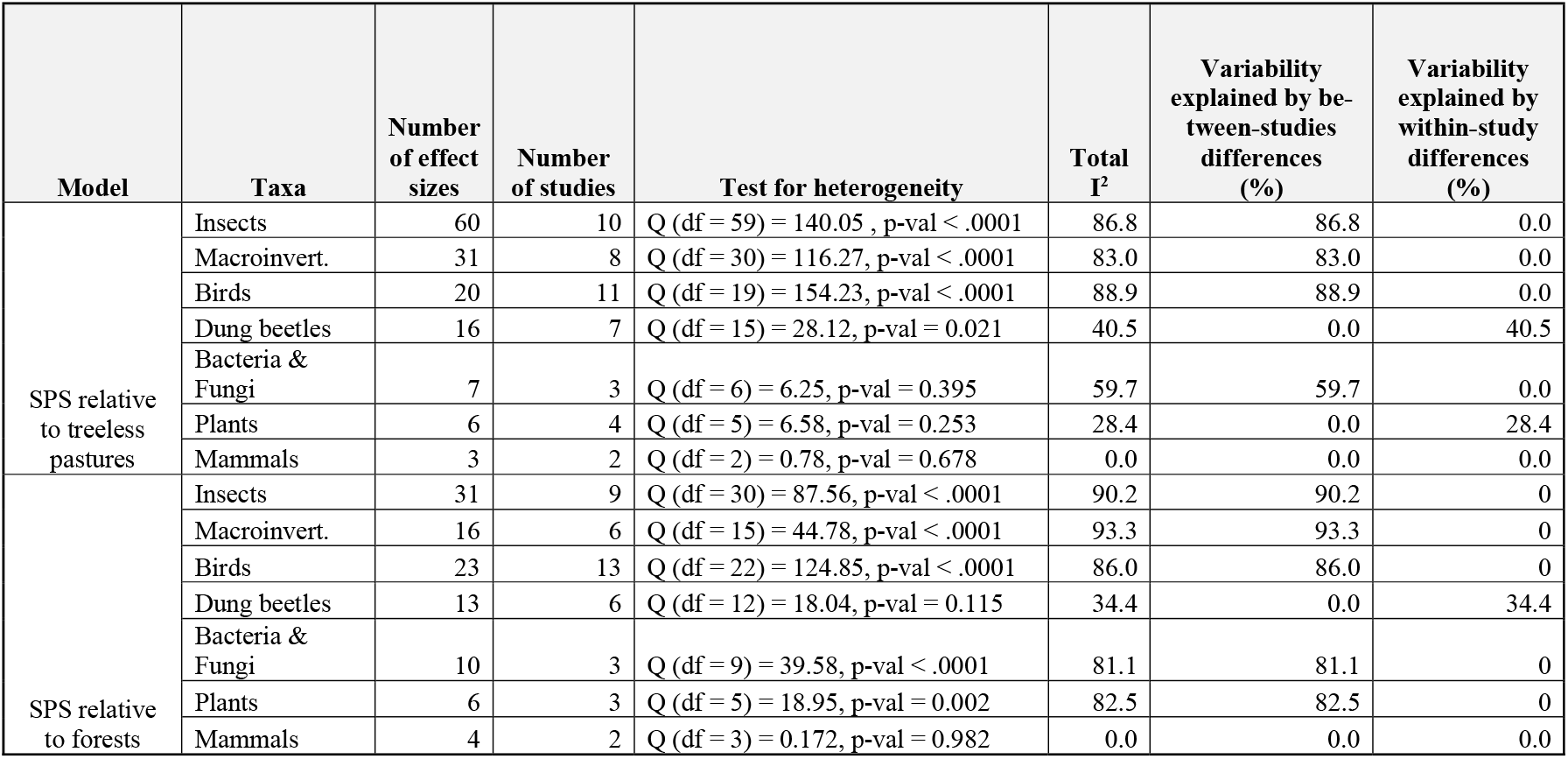
Assessment and quantification of heterogeneity in meta-analytic models across taxonomic groups. A Cochran’s heterogeneity Q-test (*78*) evaluates whether the observed effect sizes’ variability is greater than what would be expected from sampling variability alone. A significant test result indicates that true effect size differences exist between studies in our data. The I^2^ statistic shows the percentage of variation across studies that is due to heterogeneity, rather than chance (i.e., sampling error or within study variability) (*79*). I^2^ values greater than 75% indicate substantial heterogeneity (*80*).

**Table S4.** Assessment of publication and of time-lag bias in meta-analytic models across biodiversity metrics. Publication bias, which refers to the selective publication of articles finding significant effects over those finding non-significant effects, was assessed by modifying the multi-level meta-analysis model to include the square of the inverse of effective sample size as a moderator (i.e., small study effects). Biased models were identified if the slope estimate of the moderator significantly differed from zero (i.e., P < 0.05) (*72*). Time-lag bias, which occurs when statistically significant effects are published more quickly than smaller or non-statistically significant effects, was also tested by including the year of publication (mean-centered) as a moderator in the meta-analysis model. Biased models were identified if the slope estimate of the moderator differed significantly from zero (i.e., P < 0.05) (*72*). When the bias-adjusted overall effect size maintains the same sign and significance, the meta-analysis results are deemed robust against possible publication and time-lag bias. For models where there was no evidence of publication or time-lag bias (P > 0.05 for both, indicated by NA), we did not calculate the bias-adjusted effect size.

| Model | Biodiversity metric | Publication bias (P-value)* | Time-delay effects (P-value)** | Effect size mean Hedges' g (95% CI) | Effect size adjusted for publication bias*** mean Hedges' g (95% CI) |
| --- | --- | --- | --- | --- | --- |
| SPS relative to treeless pastures | Overall | <.0001 | 0.006 | 1.115 (0.615, 1.616) | 1.060 (0.564, 1.560) |
|  | Richness | 0.001 | 0.119 | 1.304 (0.652, 1.957) | 1.090 (0.499, 1.680) |
|  | Abundance | 0.047 | 0.028 | 0.924 (0.203, 1.644) | 0.771 (0.022, 1.520) |
|  | Diversity | 0.025 | 0.936 | 0.996 (0.120, 1.872) | 1.130 (0.428, 1.840) |
|  | Stability | 0.096 | 0.100 | 0.290 (-0.023, 0.603) | NA |
| SPS relative to forests | Overall | 0.766 | 0.231 | -0.122 (-0.589, 0.345) | NA |
|  | Richness | 0.542 | 0.228 | -0.289 (-0.834, 0.256) | NA |
|  | Abundance | 0.101 | 0.212 | 0.247 (-0.348, 0.842) | NA |
|  | Diversity | 0.309 | 0.798 | 0.682 (-0.409, 1.772) | NA |
|  | Stability | 0.579 | 0.365 | 0.046 (-0.125, 0.218 ) | NA |

**Table S5.** Assessment of publication and of time-lag bias in meta-analytic models across biogeographic regions. Publication bias, which refers to the selective publication of articles finding significant effects over those finding non-significant effects, was assessed by modifying the multi-level meta-analysis model to include the square of the inverse of effective sample size as a moderator (i.e., small study effects). Biased models were identified if the slope estimate of the moderator significantly differed from zero (i.e., P < 0.05) (*72*). Time-lag bias, which occurs when statistically significant effects are published more quickly than smaller or non-statistically significant effects, was also tested by including the year of publication (mean-centered) as a moderator in the meta-analysis model. Biased models were identified if the slope estimate of the moderator differed significantly from zero (i.e., P < 0.05) (*72*). When the bias-adjusted overall effect size maintains the same sign and significance, the meta-analysis results are deemed robust against possible publication and time-lag bias. For models where there was no evidence of publication or time-lag bias (P > 0.05 for both, indicated by NA), we did not calculate the bias-adjusted effect size.

| Model | Biogeographic region | Small study effects<br>(P-value) | Time-delay effects<br>(P-value) | Effect size<br>mean Hedges' g (95% CI) | Effect size adjusted for<br>publication bias<br>mean Hedges' g (95% CI) |
| --- | --- | --- | --- | --- | --- |
| SPS relative to treeless pastures | Tropics | 0.001 | 0.022 | 1.248 (0.598 -1.897) | 1.110 (0.461, 1.760) |
|  | Subtropics | 0.001 | 0.003 | 1.628 (0.413, 2.844) | 1.390 (0.714, 2.060) |
|  | Mediterranean | 0.122 | - | -0.255 (-0.603, 0.092) | NA |
|  | Temperate | 0.461 | 0.911 | 0.581 (-0.911, 2.072) | NA |
| SPS relative to forests | Tropics | 0.835 | 0.348 | -0.271 (-0.916, 0.374) | NA |
|  | Subtropics | 0.482 | 0.717 | -0.320 (-0.731, 0.091) | NA |
|  | Mediterranean | 0.118 | 0.066 | 0.644 (-0.593, 1.881) | NA |
|  | Temperate | 0.362 | 0.852 | 0.488 (-1.193, 2.170) | NA |

**Table S6.** Assessment of publication and of time-lag bias in meta-analytic models across taxonomic groups. Publication bias, which refers to the selective publication of articles finding significant effects over those finding non-significant effects, was assessed by modifying the multi-level meta-analysis model to include the square of the inverse of effective sample size as a moderator (i.e., small study effects). Biased models were identified if the slope estimate of the moderator significantly differed from zero (i.e., P < 0.05) (*72*). Time-lag bias, which occurs when statistically significant effects are published more quickly than smaller or non-statistically significant effects, was also tested by including the year of publication (mean-centered) as a moderator in the meta-analysis model. Biased models were identified if the slope estimate of the moderator differed significantly from zero (i.e., P < 0.05) (*72*). When the bias-adjusted overall effect size maintains the same sign and significance, the meta-analysis results are deemed robust against possible publication and time-lag bias. For models where there was no evidence of publication or time-lag bias (P > 0.05 for both, indicated by NA), we did not calculate the bias-adjusted effect size.

| Model | Taxa | Small study effects<br>(P-value) | Time-delay effects<br>(P-value) | Effect size<br>mean Hedges' g (95% CI) | Effect size<br>adjusted for<br>publication bias<br>mean Hedges' g (95% CI) |
| --- | --- | --- | --- | --- | --- |
| SPS relative<br>to treeless<br>pastures | Insects | 0.210 | 0.166 | 1.250 (0.084, 2.415) | NA |
|  | Macroinvert. | 0.002 | 0.065 | 1.085 (0.143, 2.026) | 1.180 (0.497, 1.850) |
|  | Birds | 0.122 | 0.208 | 1.115 (-0.079, 2.308) | NA |
|  | Dung beetles | 0.007 | 0.952 | 0.987 (0.525, 1.448) | 1.120 (0.426, 1.810) |
|  | Bacteria & Fungi | 0.832 | 0.443 | 0.095 (1.179, 1.369) | NA |
|  | Plants | 0.125 | 0.603 | 0.636 (0.172, 1.100) | NA |
|  | Mammals | 0.884 | 0.813 | 0.142 (-1.046, 1.330) | NA |
| SPS relative<br>to forests | Insects | 0.539 | 0.791 | 0.200 (-0.845, 1.245) | NA |
|  | Macroinvert. | 0.761 | 0.693 | -0.096 (-1.723, 1.530) | NA |
|  | Birds | 0.833 | 0.189 | -0.125 (-0.863, 0.611) | NA |
|  | Dung beetles | 0.073 | 0.583 | -0.6762 (-1.158, -0.195) | NA |
|  | Bacteria & Fungi | 0.849 | 0.978 | 0.742 (-1.028, 2.512) | NA |
|  | Plants | 0.923 | 0.149 | -0.313 (-2.088, 1.463) | NA |
|  | Mammals | 0.999 | 0.999 | -0.0453 (-0.647, 0.557) | NA |

## Data file S1. (separate file .xls)

### Dataset. Overview of the studies included in the meta-analyses

In addition to providing this metadata, we fully intend to make the actual data public upon the publication of this manuscript.

